# Development of properly-polarized trophoblast stem cell-derived organoids to model early human pregnancy

**DOI:** 10.1101/2023.09.30.560327

**Authors:** J Zhou, MA Sheridan, Y Tian, KJ Dahlgren, M Messler, T Peng, T Ezashi, LC Schulz, BD Ulery, RM Roberts, DJ Schust

## Abstract

The development of human trophoblast stem cells (hTSC) and stem cell-derived trophoblast organoids has enabled investigation of placental physiology and disease and early maternal-fetal interactions during a stage of human pregnancy that previously had been severely restricted. A key shortcoming in existing trophoblast organoid methodologies is the non-physiologic position of the syncytiotrophoblast (STB) within the inner portion of the organoid, which neither recapitulates placental villous morphology *in vivo* nor allows for facile modeling of STB exposure to the endometrium or the contents of the intervillous space. Here we have successfully established properly-polarized human trophoblast stem cell (hTSC)-sourced organoids with STB forming on the surface of the organoid. These organoids can also be induced to give rise to the extravillous trophoblast (EVT) lineage with HLA-G^+^ migratory cells that invade into an extracellular matrix-based hydrogel. Compared to previous hTSC organoid methods, organoids created by this method more closely mimic the architecture of the developing human placenta and provide a novel platform to study normal and abnormal human placental development and to model exposures to pharmaceuticals, pathogens and environmental insults.

**Motivation:** Human placental organoids have been generated to mimic physiological cell-cell interactions. However, those published models derived from human trophoblast stem cells (hTSCs) or placental villi display a non-physiologic “inside-out” morphology. *In vivo*, the placental villi have an outer layer of syncytialized cells that are in direct contact with maternal blood, acting as a conduit for gas and nutrient exchange, and an inner layer of progenitor, single cytotrophoblast cells that fuse to create the syncytiotrophoblast layer. Existing “inside-out” models put the cytotrophoblast cells in contact with culture media and substrate, making physiologic interactions between syncytiotrophoblast and other cells/tissues and normal and pathogenic exposures coming from maternal blood difficult to model. The goal of this study was to develop an hTSC-derived 3-D human trophoblast organoid model that positions the syncytiotrophoblast layer on the outside of the multicellular organoid.

**Graphical abstract:** 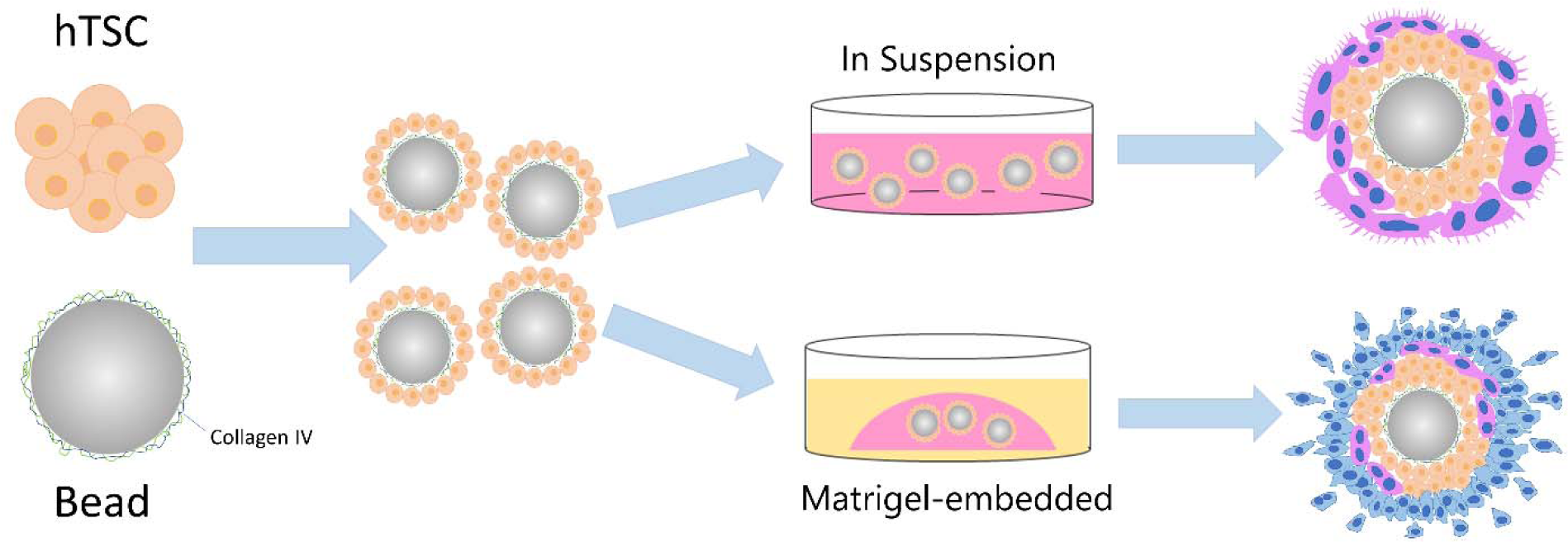

## Introduction

Trophoblast is a specialized, conceptus-derived cell type of the human placenta. All of the trophoblast lineages are generally thought to arise from the blastocyst trophectoderm (TE) [1–3], and their coordinated proliferation and differentiation is essential for a successful pregnancy [4]. Initially, some of the trophectoderm cells merge to form a leading edge pre-villous syncytial mass that implants into the endometrium. Subsequently, the three major trophoblast subpopulations of the villous stage placenta are established and maintained throughout pregnancy: cytotrophoblast cells (CTB), extravillous trophoblast cells (EVT), and syncytiotrophoblast (STB), although there may be multiple functional sub-types within these larger groupings [5]. The underlying CTB are a proliferative population that can give rise to either the STB lineage that lines the outer surface of the villi, or the EVT lineage at the point where some villous tips anchor to the maternal decidua (anchoring villi) [6–9]. Impaired trophoblast development and function are thought to lead to pregnancy complications, including miscarriage, preeclampsia, and intrauterine growth restriction [10–12].

Efforts to understand the human placenta have been plagued by a lack of functional experimental models [13]. Our current understanding of human placental morphology during early pregnancy relies upon rare, archived samples and non-human primate models. As a consequence, fundamental questions about the initial stages of human trophoblast invasion and placentation remain unanswered. Historically, trophoblast isolated from human placentas have proven to be short-lived in culture and to rapidly differentiate [14]. This limitation was recently resolved by the isolation and derivation of trophoblast stem cells [15] and the development of trophoblast organoids [16].

Several groups have reported using stem cells or first-trimester placental cells to establish three-dimensional (3D) placental organoids that contain both STB and EVT [16–19]. While these 3D organoids are thought to functionally model the human placenta, their polarity is inverted. This “inside-out” morphology, which is likely a consequence of the Matrigel droplet in which they are embedded serving as a basement membrane, results in an outer layer of CTB cells and formation of multi-nucleated STB within the center of the organoid. The goal of the present work has been to develop a method to invert this polarity, creating human trophoblast organoids from trophoblast stem cells that more closely resemble the *in vivo* structural organization of the human placenta.

## Results

### Generation of properly-polarized STB trophoblast organoids from hTSC that are characterized by an outer layer of STB

CT27 hTS cells (female) maintained in 2D culture were dispersed and cultured with collagen IV-coated beads for 2 days. The bead-associated trophoblast organoids were expanded in trophoblast organoid medium (TOM)[16] in the absence of forskolin to induce syncytialization, with organoids reaching an average size of 600 µm by day 10 (Fig. 1A). These organoids could be passaged and they regenerated their structures within 7-10 days (Fig 1A). To evaluate the morphology of the trophoblast organoids after 10 days in culture, they were fixed, paraffin-embedded, sectioned and stained with hematoxylin and eosin (H&E). The trophoblast organoids growing around a bead were composed of multiple inner layers of mononuclear cells surrounded by a semi-continuous multinucleated layer (Fig 1B, left panel), recapitulating the organization of placental villi *in vivo*. These bead-associated, properly-polarized organoids contained numerous vacuoles, also typical of the STB *in vivo* [20–22] and during organ culture [23, 24]. When grown under the same conditions in the absence of the collagen-coated bead, however, these same multinucleated cells (STB) were located almost entirely in the center of the organoids, and the outer layers of the organoid harbored multiple rows of mononucleated cells (Fig. 1B, right panel). We have replicated these methods using a second, male hTSC line, CT29, with resulting generation of organoids that also demonstrate a properly-polarized morphology (data not shown).

**Fig. 1:**
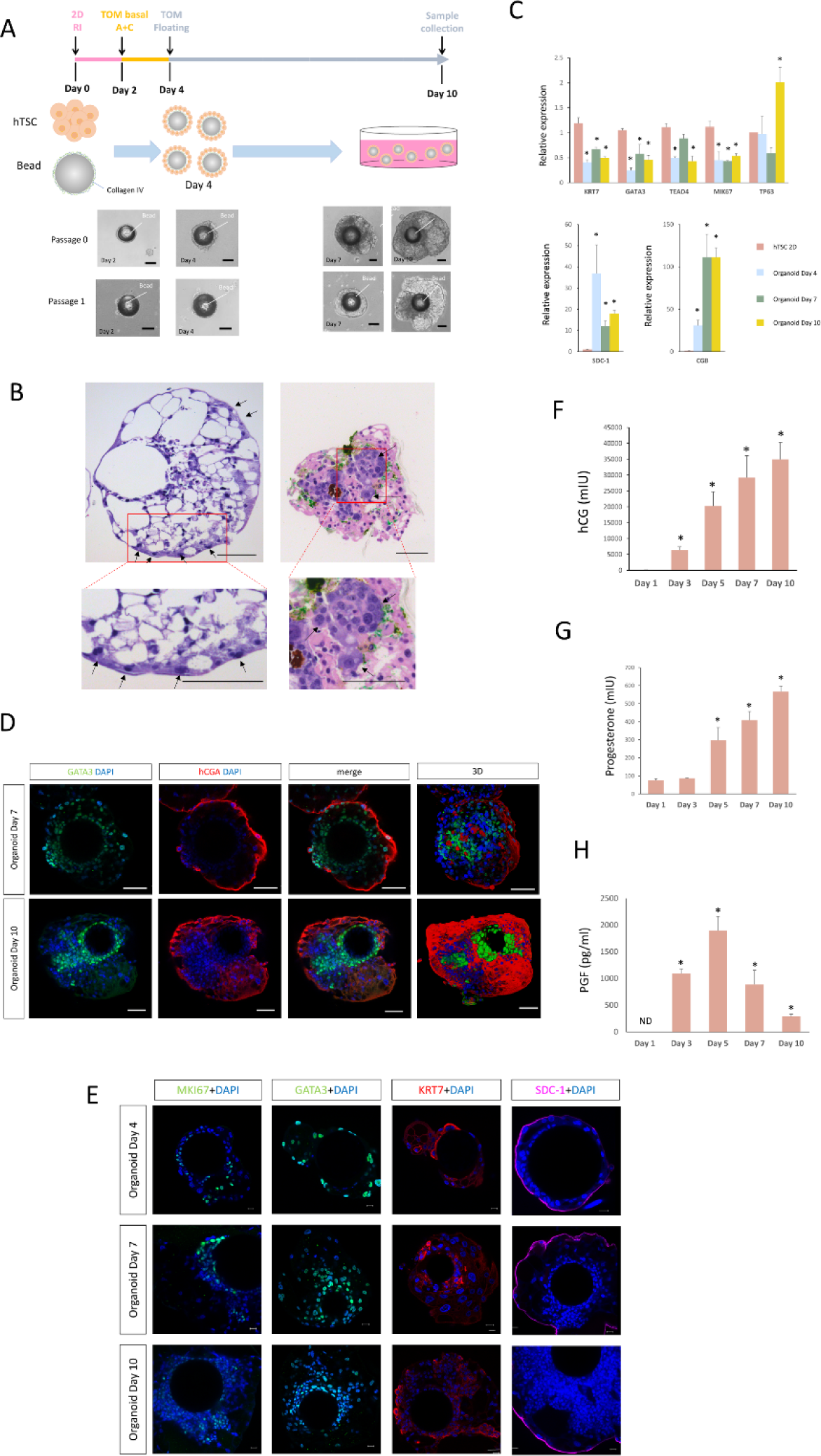
Establishment of properly-polarized trophoblast organoid cultures from hTSC cells. **A.** The hTSC cell line (CT27) was maintained under traditional 2D-culture conditions. To develop our organoid cultures, these cells were dispersed and incubated with collagen IV-coated beads cultured in TSC medium supplemented with 5 µM Y27623 for 2 days (pink line). The medium was then switched to TOM basal medium containing A83-01(500 nM) and CHIR (1.5 µM) (TOM basal A+C) for an additional 2 days (yellow line). The developing organoids were then treated with complete TOM medium until day 10 (grey line). Bright-field images of hTSC-derived trophoblast organoids from passage 0 and passage 1 are depicted. Scale bars, 200µm. **B.** Hematoxylin and eosin (H&E) staining of the organoids on day 10 revealed multinucleated structures (black arrows) on the surface of the organoid in the presence of the bead (left panel); the multinucleated areas (black arrows) remained inside the organoid in absence of the bead (right panel). Scale bars, 50µm. **C.** Quantitative real-time PCR analysis of CTB markers (upper panel) and STB markers (lower panel) in hTSC (2D culture) and STB organoids on Day 4, Day 7 and Day 10. *KRT7*, *GATA3*, *TEAD4*, and *MKI67* transcript levels were higher in hTSC than in organoids, while *hCGB* and *SDC1* transcript levels were much lower in hTSC than in organoids. The data were normalized to *GAPDH* expression. Error bars indicate SDs (n=3). **D.** Immunostaining of hCGA and GATA3 in STB organoids on Day 7 and Day 10. hCGA is expressed in multinucleated areas lining the outer surface of the organoids while the mononucleated cells closer to the center of the organoids are GATA3 positive. Reconstructed 3D confocal images showing the distribution of the hCGA and GATA3. **E.** Immunostaining of MKI67, GATA3, KRT7, and SDC-1 in trophoblast organoids on Day 4, Day 7 and Day 10. Trophoblast cells in the center of the organoids express the CTB markers GATA3 and KRT7; MKI67 staining labeled proliferating trophoblast cells; and the STB marker SDC-1 is detected on the surface of the trophoblast organoids. Images are representative of three biological replicates. Daily production of three placental hormones hCG **(F)**, progesterone **(G)**, and PGF **(H)** was assessed in the culture medium by ELISA over the time course of organoid culture. The medium was replaced on day 2, day 4, day 6 and day 9 of culture, 24h before collection. Presented values are the means ± SDs for three separate experiments.

To further characterize our properly-polarized trophoblast organoids, quantitative PCR (qPCR) and immunofluorescent staining was performed on organoids collected on culture days 4, 7 and 10. Using qPCR, we detected significantly higher levels of *SDC-1* (Syndecan 1) and *CGB* (chorionic gonadotropin beta) transcripts in trophoblast organoids than in undifferentiated hTSCs grown in a two-dimensional configuration in tissue culture wells (Fig. 1C). Expression of the STB markers, SDC-1, CGB and CGA was detected mainly in multinucleated patches near the outside surface of the organoids, antigens such as MKI67 (marker of proliferation Ki-67), GATA3 (GATA binding protein 3) and KRT7 (cytokeratin-7), were largely concentrated in the mononucleated cells inside the organoids (Fig 1 D. E, Fig 3 A-B).

The trophoblast organoids secrete hCG into the medium, which is detectable after 1 day in culture and gradually increases until day 10 (Fig. 1F), whereas hTSC-derived organoids without beads and maintained under the conditions described by Sheridan et al. [25] produced hCG very poorly. Like hCG, progesterone (P4) release from bead-associated properly oriented organoids also increased over time (Fig. 1G). Production of placental growth factor (PGF) peaked at day 5 in culture and dropped significantly by day 10 (Fig. 1H).

### Ultrastructure of properly polarized trophoblast organoids

Scanning electron microscopy (SEM) and transmission electron microscopy (TEM) were used to visualize common structural features of trophoblast organoids and *in vivo* STB [16]. The trophoblast organoids contained multinucleated areas surrounded by a continuous cell border and displayed large areas of surface microvilli consistent with normal villous STB structure *in vivo* (Fig 2A, 2D). The internal mononucleated cells in the organoids displayed irregularly shaped nuclei, with chromatin aggregation beneath the nuclear membrane. The multinucleated cells plasma membrane tended to be irregular with microvilli projections (Fig 2B, 2D). Mononuclear cells maintained contact with each other through adhesion complexes, including tight junctions, adherens junctions and desmosomes (Fig 2B). We also noted that the mitochondria found in multinuclear patches (STB) have a distinct appearance. Similar to prior reports on placental biopsies [26], the mitochondria in the multinuclear patches were numerous, round, electron-dense, and contained visible tubular cristae; they were also smaller than those found in the mononuclear cells inside the organoids (Fig 2C, 2E). The reduction in mitochondrial size and structural alterations in mitochondrial cristae are consistent with the decrease of glycolysis and expansion of steroidogenic activity typically seen in STB *in vivo* [27, 28]. Also as previously reported[29], numerous autolysosomes and autophagosomes were detected in both organoid cell types (STB and CTB) (Fig 2B, 2D), suggesting a potential to activate autophagy, which is involved in trophoblast fusion [30]. We also located a non-membrane-bound cytoplasmic organelle, termed a nematosome, in the cytoplasm of a trophoblast organoid cell (Fig 2E). Nematosomes have previously been reported in human trophoblast, but their function remains unclear [31].

**Fig. 2:**
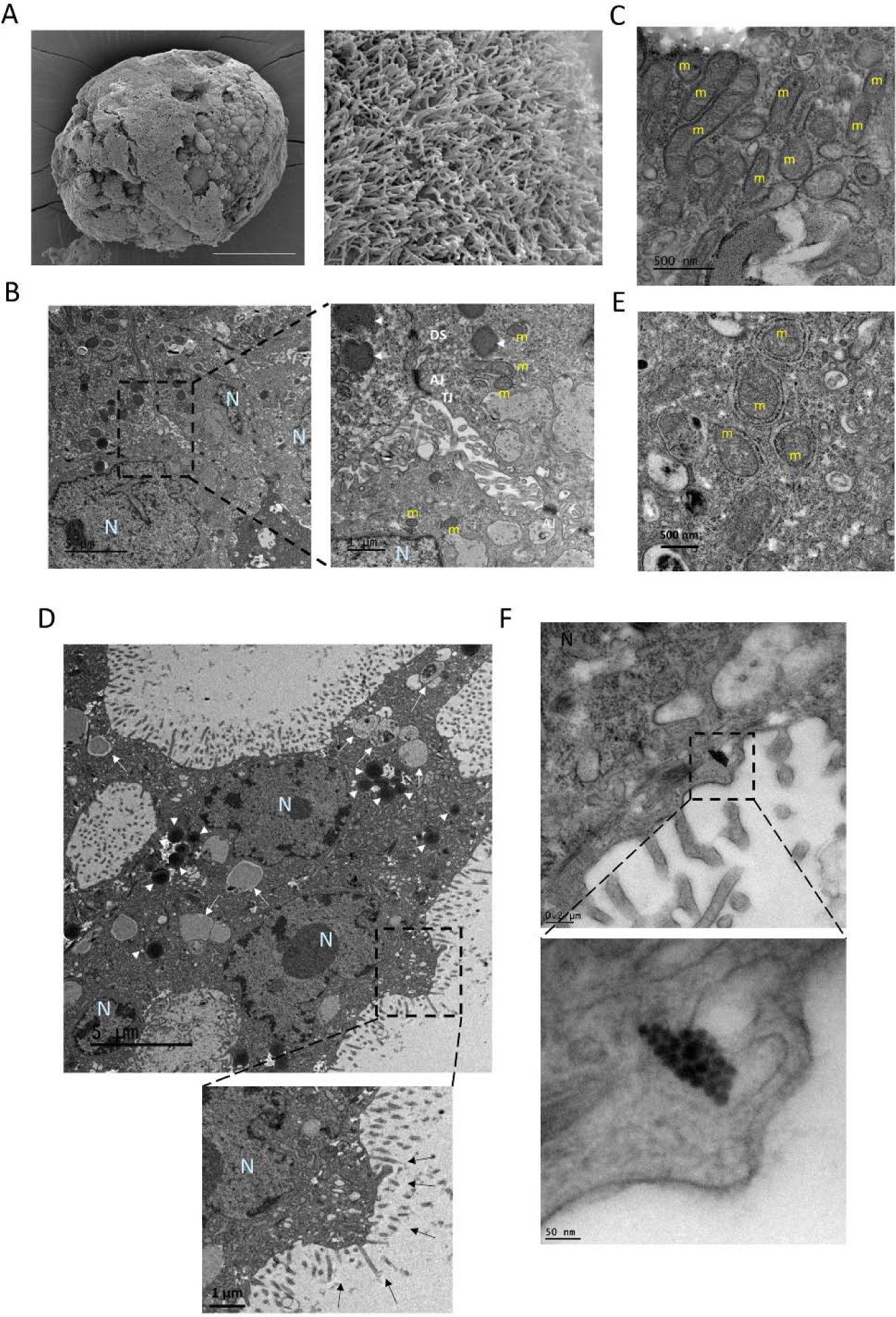
Ultrastructure of CTB and STB in trophoblast organoids. A. Scanning electron microscopy (SEM) images of STB organoids reveals a topography (left panel, scar bar, 100 µm) covered by dense microvilli (right panel, scar bar, 3 µm). **B.** Transmission electron microscopy (TEM) images of an “apical junctional complex” of mononuclear CTB from 3D floating STB-covered organoids. Depicted are the tight junctions (TJ), directly beneath the microvilli; adherens junctions (AJ), and desmosomes (DS) below the apical junctional complex. Scare bars, 5 µm and 1 µm. N, Nucleus; m, mitochondria; lysosome (white arrow heads). **C.** The larger mitochondria in presumed mononuclear CTB have lamellar cristae. Scare bar, 500 nm. **D**. Microvilli (black arrow), lysosomes (white arrow heads) and autolysosomes (white arrow) that can be seen in multinuclear STB at Day 10. Scale bars, 5 µm and 1 µm. **E.** STB mitochondria have a dense matrix and vesicular cristae. Scale bar, 500 nm. **F.** A nematosome can be seen in a mononuclear trophoblast cell. Scar bars, 200 nm, 50 nm.

### HLA-G+CGB+ double positive cells were detected in properly polarized trophoblast organoids

It has been reported that a few HLA-G^+^ cells can be found in inverted human trophoblast organoids maintained in a standard proliferation medium TOM [16]. We also observed sporadic cells in the inner layer of our trophoblast organoids that expressed the EVT marker, HLA-G, on Days 4, 7, and 10 (Fig. 3A-B). In addition, we occasionally found HLA-G^+^ CGB^+^ SDC-1^+^ cells located on the surface of the properly polarized floating organoids; these individual cells could also be retrieved from collected culture media (Fig. 3C, 3D).

**Fig. 3:**
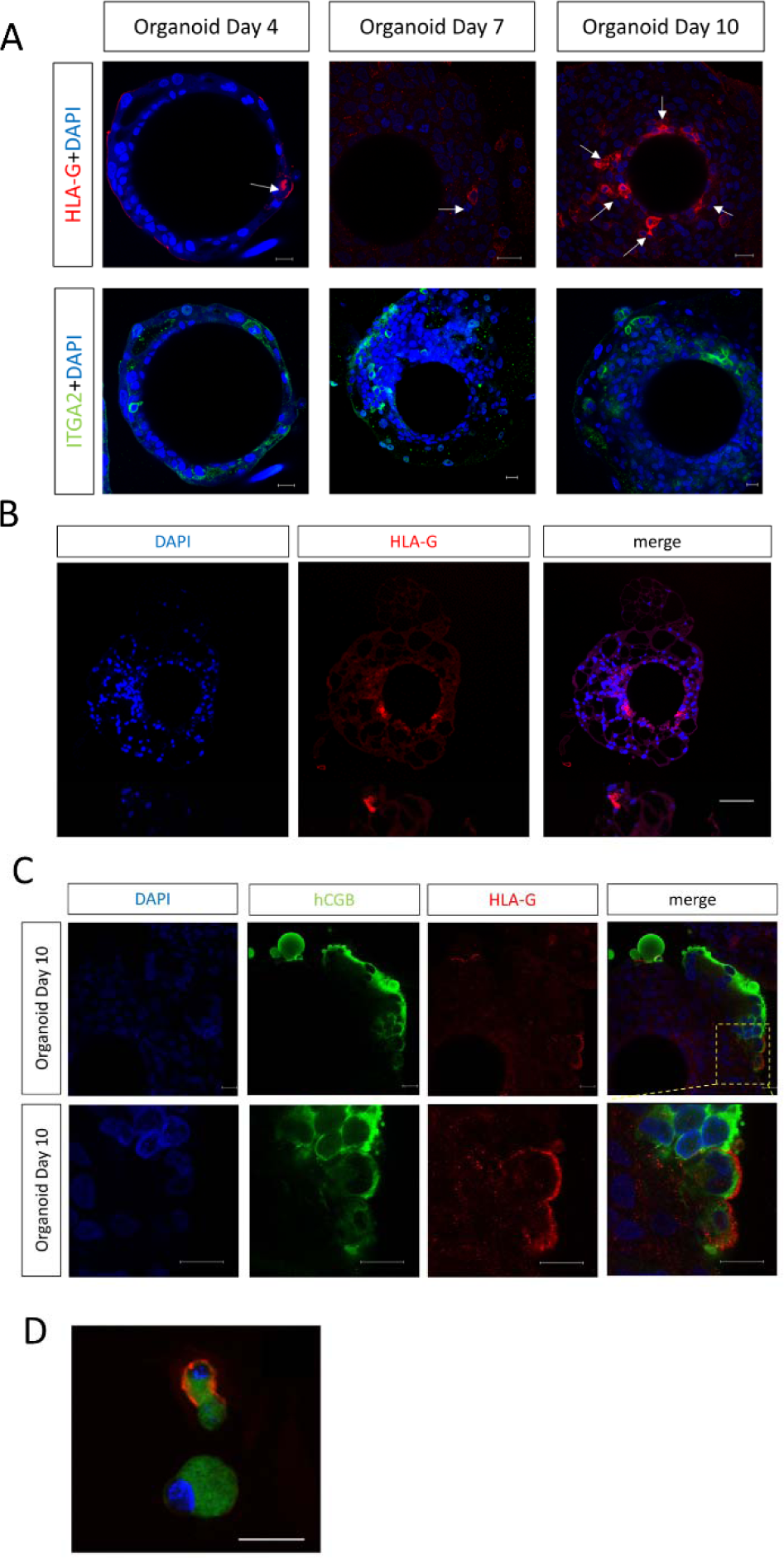
Expression of HLA-G in trophoblast organoids. **A.** Whole mount staining shows sporadic expression of the EVT markers, HLA-G and ITGA2, in floating STB organoids on Day 4, Day 7 and Day 10. Scare bars, 20 µm. **B.** Sections of paraffin-embedded STB organoids at Day 10 were immunostained for HLA-G. A small number of cells close to the beads were HLA-G positive. Scare bars, 100 µm. **C.** Co-staining of HLA-G and hCGB using whole-mounted floating STB organoids with images showing HLA-G staining on hCGB positive cells. Scare bars, 20 µm. **D.** Cells shed into the culture medium co-expressed HLA-G and hCGB. Scare bar, 20 µm.

### Generation of migratory and invasive EVT-like cells from properly-polarized trophoblast organoids

Two-dimensional cultures of hTSC and human trophoblast organoids can be coaxed to give rise to EVT-like cells by using specialized media (EVTM) that contains a TGFβ inhibitor and neuroregulin 1 (NRG1) [15, 16, 32]. We adapted this EVTM protocol for our trophoblast organoids to induce similar differentiation. After 7 days of floating culture in suspension medium (Fig. 1A), we transferred the trophoblast organoids into Matrigel droplets and switched to culture in EVTM (Fig. 4A). Cells were observed to have migrated outside of the organoid into the Matrigel within 7 days, forming spindle-like structures with typical EVT morphology, as reported by others [33–35] (Fig. 4E-F). H&E-stained sections of the organoids grown in Matrigel and EVTM showed heterogeneous cell populations (Fig 4B). Mononuclear cells containing nuclei of various size and multinucleated patches enclosed by large areas of cytoplasm could be identified. Many of the cells in these organoids at day 14 (7 days in Matrigel) expressed both the CTB column marker ITGA2 (Integrin α2) and the EVT marker HLA-G [36], with the most intense HLA-G staining located on the outermost layer of the organoid (Fig. 4C). We identified proliferating cells by staining for MKI67, which is expressed during all cell cycle stages except G0 [37, 38]. Most MKI67+ cells within the organoids cultured in EVTM for 7 days were mononuclear and contained nuclei of consistent size; these were distributed throughout the organoid (Fig. 4D). Note that at the time when the trophoblast organoids were initially seeded into the Matrigel in EVTM, they were already surrounded by multinucleated STB and only rare cells expressed ITGA2 (Fig. 1E and 3A). After 7 days in EVTM, ITGA2 was present in a majority of the mononucleated cells, suggesting that the CTB columns formed and developed in response to Matrigel and EVTM conditions (Fig. 4C). We term these organoids EVT organoids.

**Fig. 4:**
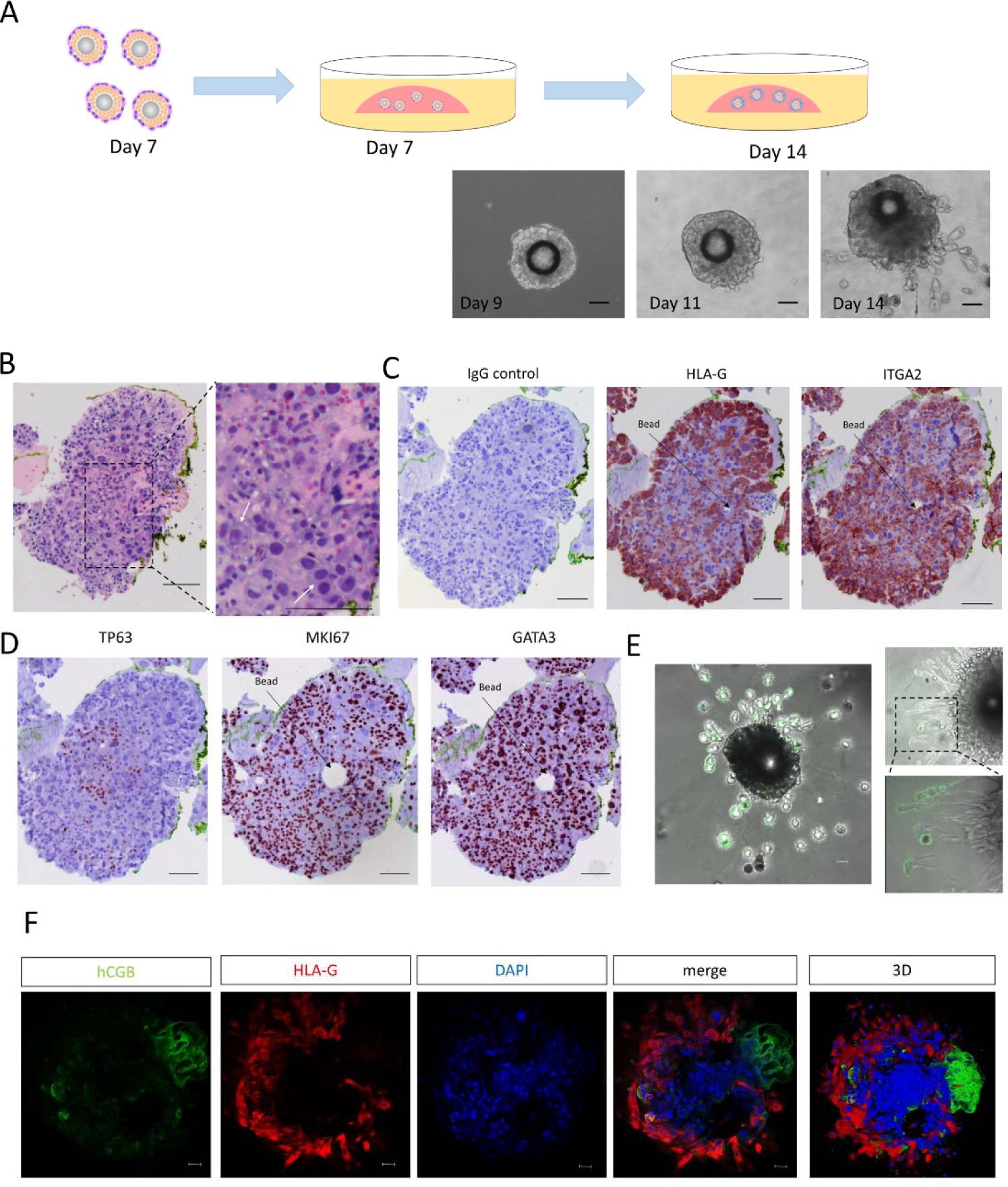
Generation of migratory and invasive HLA-G^+^ EVT-like cells from trophoblast organoids. **A.** STB organoids on Day 7 were plated into Matrigel in EVT differentiation medium (EVTM). Phase-contrast images on Day 9, Day 11 and Day 14 (2 days, 4 days and 7 days after placement into the Matrigel drops, respectively). Scale bars, 200µm. **B.** H&E staining displayed large cells (compared with the adjacent trophoblast cells) containing two nuclei enclosed in a voluminous cytoplasm (white arrows). Paraffin-embedded sections of organoids in EVTM at Day 14 stained for the EVT markers, HLA-G and ITGA2 (**C**); and the CTB markers TP63, MKI67 and GATA3 (**D**). Scare bars, 100 µm. **E.** Live cell imaging showing cells growing out from organoids in Matrigel and EVTM stained with the anti-HLA-G monoclonal antibody G233. Cells stream out of organoids in two ways: migrating into the Matrigel (E) or adhering to and moving along the plastic dish (F). Scale bar, 50 µm. **F.** At Day 14, organoids in EVTM were co-stained for HLA-G and hCGB. hCGB is expressed at the surface of the organoids while the migrated and invaded cells from the organoid are HLA-G positive. Scale bars, 50 µm.

## Discussion

Despite its critical role in pregnancy, the placenta is the least understood human organ[39], largely a consequence of practical, legal and ethical restrictions on studying its development. Recently, several groups have reported the development of trophoblast organoids derived from first-trimester placental tissues, term placental tissues, hTSC, and naïve hPSC (human pluripotent stem cells) [16, 17, 40, 41]. Trophoblast organoids (TO) are self-renewing, three dimensional cellular structures generated *in vitro* that recapitulate multiple aspects of placenta development and function[16, 17]. However, each of the reported trophoblast organoid models to date poorly recapitulates the anatomy of the developing primitive and villous human placenta, each of which is covered by a leading edge of implanting embryo or villous surface in placenta of multinucleated syncytialized trophoblast. This STB layer has several functions, including serving as: 1) the major transporting epithelium in the placenta, with capacity for polarized gas and nutrient transfer[42], 2) a physical and immunologically-active barrier to fetal infection and toxin exposures, and 3) an induction site for immunotolerance of the antigenically foreign fetus in normal pregnancies[43, 44] through the shedding syncytial nuclear aggregates and exosomes that interact with the maternal immune system. Here we present an improved trophoblast organoid model derived from hTSC in the absence of forskolin exposure that presents its multinuclear layer at the outer surface of the organoids, more closely mimicking the *in vivo* architecture of the implanting embryo and villous placenta than prior models.

The human blastocyst is a roughly spherical structure with a diameter of about 200 µm at approximately 7 days post fertilization [45] when it undergoes implantation [46]. Size equivalent beads were modified via amidation to offer opportunities for covalent coupling with collagen type IV[47], which can provide structural support regulating adhesion, migration and survival of cells [48], and has been reported to affect the invasive behavior of trophoblast cells at the fetal-maternal interface[49]. CT27 hTSCs were exposed to amidated, collagen IV-coated beads and then cultured in suspension medium. By day 4, the developing organoids are multilayered with a syncytialized surface layer beginning to form and an inner core of proliferative single CTB-like cells that are capable of further differentiation (Fig. 1D). By day 7-10 of culture, a layer of cells with ultrastructural characteristics of intermediate stage trophoblastas described by Jones and Fox[50] can be detected between the undifferentiated CTB cells near the bead and the outer syncytium (Fig. 2B) [51–54].

The human placenta is an endocrine organ, generating a wide array of hormones required for the establishment and maintenance of pregnancy. We explored the secretory activity of properly polarized trophoblast organoids. Typical of STB *in vivo*, the properly-polarized trophoblast organoids secrete hCG, P4 and PGF into the surrounding culture media, although secretion of the latter occurs over a limited time and does not occur until a later time point in the culture period. It is uncertain why PGF secretion exhibits this pattern in our organoid cultures; one possible explanation is the absence of stimuli such as hypoxia, inflammatory cytokines, growth factors and hormones normally present during implantation and placental development *in vivo* [55–58]. Our properly-oriented trophoblast organoids contain vacuoles similar to those seen in the inverted 3D trophoblast organoids reported by Turco et al[16], grown in a medium similar to ours, and by Okae et al [15] grown in medium containing forskolin. The formation of these lumen-like glandular structures may reflect the enhanced secretory ability of intermediate-stage trophoblast in the process of differentiating from CTB to either the STB or EVT linage.

The STB of the human placental barrier is covered with microvilli that are exposed to the maternal blood in the intervillous space[59]. This microvillous surface is vital for placental nutrient exchange, transfer of oxygen, immune protection, excretion of waste products and secretion of hormones [60, 61]. Microvilli are also involved in the first intercellular interactions between mother and embryo during implantation, maintaining close adhesion between the trophoblast surrounding the implanting blastocyst and the uterine epithelium [62]. The syncytialized outer layer of our trophoblast organoids exhibit well-organized apical microvilli characteristic of the STB surface *in vivo* [63] [64] (Fig. 2A). The sporadic HLA-G^+^ CGB^+^ SDC-1^+^ cells on the surface of our organoids or retrieved from collected trophoblast organoid culture media (Fig. 3C-D) may perhaps model the remarkable and unique previllous invasive primitive STB. Primitive STB secretes massive amounts of hCG to rescue the maternal corpus luteum and thereby maintain pregnancy but also migrates into the maternal decidua to establish initial interactions with the decidual stromal cells and glands as well as decidual immune cells, including maternal dNKs [65, 66]. This hypothesis will be investigated in future experiments but is consistent with data from extended *in vitro* human blastocyst culture (34).

During the first and second trimesters of pregnancy, the CTB, villous trophoblastic columns, intermediate trophoblast and multinucleated intermediate trophoblast cells strongly express GATA3 *in vivo* while the STB are GATA3 negative [67]. The villous CTB marker, TP63 (tumor protein 63), is essential for maintaining CTB stemness and is therefore only expressed in proliferative CTB [68]; TP63 expression has also been shown to inhibit EVT migration [69]. In our study, EVT induction via exposure to a Matrigel substrate and EVT media resulted in decreased expression of TP63, while GATA3 remained highly expressed in most of the EVT-like cells (Fig 4D), which appear to penetrate the STB layer and migrate into the Matrigel (Fig 4A, 4F). Our data is consistent with the hypothesis that the column CTB ultimately give rise to the villous placenta[35]. The other proliferative cells reported to be present in the first trimester human placenta, which are MKI67^+^TP63^−ITGA2+^, appear to retain their proliferative capacity [70] and are thought to be a source of trophoblast progenitor cells located within the cell columns (Fig. 4C-D) [36]. Staining patterns for these markers in our trophoblast organoids likely reflect these trophoblast lineages and their development over time within the floating and Matrigel-embedded culture conditions.

In summary, we describe here a method to reliably generate human trophoblast organoids with a structure that more closely mimics the architecture of the human implanting embryo *in vivo* than prior trophoblast organoid models. This *in vitro* model should improve our ability to study placental susceptibility to and vertical transmission of pathogens, maternal-fetal interactions across the villous placenta, and the effects of exposure to maternal blood-borne environmental toxins and pharmaceuticals in a physiologically relevant way.

### Limitation of the study

The properly polarized model presented in this study was derived from hTSCs. Although we have replicated our results in two separate hTSC lines (one male and one female), we have not demonstrated a broader application of the method by using 1^st^-trimester primary trophoblast cells or term trophoblast cells. The properly-polarized organoids have only been expanded and passaged a limited number of times in order to achieve optimized *in vitro* culture conditions. Future experiments will test the maximum number of passages and *in vitro* culture duration that these organoids can tolerate. The organoids generated here are comprised of trophoblast derivatives but not other fetal cells present within the placental villi, including fetal mesenhchymal and endothelial cells or maternal cells, including decidual cells and immune cells, normally found at the site of implantation *in vivo*. This limitation will be addressed in future experiments using co-culture models. The conditions described here generate mixed cultures of bead-attached, properly-polarized organoids mixed with standard inverted trophoblast organoids that develop in the absence of beads; the two organoid types must be separated to specifically study properly-polarized 3D development. Finally, like many TSC-derived models, the properly-polarized trophoblast organoids are comprised of several cell types and exhibit some degree of heterogeneity among individual organoids in term of size and number of cells. Therefore, these organoids will require dispersion and single cell and/or single nuclei analyses for some applications.

## STAR METHODS

### KEY RESOURCES TABLE

| REAGENT or RESOURCE | SOURCE | IDENTIFIER |
| --- | --- | --- |
| Antibodies |  |  |
| SDC-1 | NOVUS | AF2780 |
| KRT7 | Dako | M7018 |
| hCGB | Abcam | Ab9582 |
| GATA-3 | Cell Signaling | 5852S |
| hCGA | NOVUS | NBP2-29428 |
| TP63 | Cell Signaling | 13109S |
| HLA-G | Santa Cruz | sc-2179 |
| HLA-G (FITC) | NOVUS | NB110-55298F |
| MKI67 | Abcam | ab66667 |
| ITGA2 | Abcam | ab181548 |
| Alexa Flour 488 Donkey anti-Rabbit IgG | Invitrogen | A21206 |
| Alexa Fluor 555 donkey anti-Mouse IgG | Invitrogen | A-31570 |
| Alexa Fluor 647 donkey anti-Goat IgG | Invitrogen | A-21447 |
| Chemicals, peptides, and recombinant proteins |  |  |
| Sulfo-NHS | Thermo Scientific | PG82071 |
| 1-(3-dimethylaminopropyl)-3-ethylcarbodiimide hydrochloride | Thermo Scientific | 171440010 |
| sodium bicarbonate | Sigma-Aldrich | S5761 |
| ethylenediamine | Sigma-Aldrich | E26266 |
| Collagen IV, mouse | Corning | 354233 |
| Matrigel™ GFR Membrane Matrix | Corning | 356230 |
| Dimethyl Sulfoxide | Sigma-Aldrich | D8418 |
| TrypLE Express Enzyme (1X), no phenol red | Gibco | 12604-021 |
| Fetal Bovine Serum, embryonic stem cell-qualified, US origin | Gibco | 16141061 |
| BSA (Fatty Acid Free) | Fisher Scientific | BP9704100 |
| ITS-X | Gibco | 51500056 |
| KnockOut Serum Replacement | Gibco | 10828010 |
| hEGF (2D culture) | Sigma | E9644-.2MG |
| Stemolecule™ SB431542 | Reprocell USA | 04-0010 |
| Y27632 | Reprocell USA | 04-0012-02 |
| CHIR99021 | Reprocell USA | 04-0004 |
| A83-01 | Reprocell USA | 04-0014 |
| 2-mercaptoethanol | Sigma | 21985-023 |
| L-Ascorbic acid 2-phosphate sesquimagnesium salt hydrate | Sigma | A8960-5G |
| Valproic acid sodium salt | Sigma | P4543-10G |
| Human Neuregulin-1 (hNRG-1) | Cell Signaling | 26941S |
| Forskolin | Sigma | F6886-10MG |
| N2 supplement | Gibco | 17502048 |
| B27 supplement minus vitamin A | Gibco | 12587010 |
| N-acetyl-L-cysteine | Sigma | A9165-5G |
| Recombinant human EGF (3D culture) | Peptotech | AF-100-15 |
| Recombinant human R-spondin 1 | BioTechne | 4645-RS |
| GlutaMAX | Gibco | 35050061 |
| Recombinant human FGF2 | Peptotech | 100-18C |
| PGE2 | Sigma | P0409 |
| Recombinant human HGF | Peptotech | 100-39 |
| donkey serum | Sigma | D9663 |
| BSA (immunofluorescence) | Sigma | A7906 |
| Triton X-100 | Sigma | X100 |
| DAPI | Invitrogen | D3571 |
| VECTASHIELD Mounting Medium | VectorLabs | H-1200 |
| formalin | Electron Microscopy Sciences | 15740-01 |
| PFA | Electron Microscopy Sciences | 15710 |
| osmium tetroxide | Ted Pella, Inc. | 18459 |
| Critical commercial assays |  |  |
| Vectastain Elite ABC-HRP kit | Vector Lab | PK-6100 |
| hCG ELISA kits | Genway bio | GWB-BQK0F2 |
| P4 ELISA kits | Genway bio | GWB-BQK0FC |
| PGF human ELISA kits | Invitrogen | EHPGF |
| RNeasy Mini kit | QIAGEN | 74104 |
| cDNA synthesis kit | BIO-RAD | 1708890 |
| SYBR Green PCR | MedChemExpress | HY-K0501 |
| TURBO-DNA-free kit | Invitrogen | AM1907 |
| Experimental models: Cell lines |  |  |
| hTSC CT27 | Okae et al. 2018 | N/A |
| hTSC CT29 | Okae et al. 2018 | N/A |
| Software and algorithms |  |  |
| Prism 5.0 | GraphPad | www.graphpad.com/ |
| LAS X | Leica | www.leica-microsystems.com/products/microscope-software/p/leica-las-x-ls/downloads/ |
| Other |  |  |
| polystyrene beads | Phosphorex | 170 |
| round-bottom test tube | Corning | 352059 |
| Tissue culture test plates 6-well | TPP | 92006 |
| Corning™ Costar™ Ultra-Low Attachment | Corning | 3471 |
| Microplates 6-well |  |  |
| Tissue culture dish 34mm | TPP | 93040 |
| 35 mm Dish with glass coverslip | MatTek | P35G-1.5-14-C-GRD |

### Resource Availability

#### Lead contact

Information and request for resources and reagents can be directed to and will be fulfilled by the Lead Contact, Danny J. Schust, MD at

#### Materials availability

This study did not generate new or unique reagents.

### Experimental model and subject details

No new experimental animal models or cell lines were generated for this study or were research subjects enrolled.

### Method details

#### Preparation of Beads

Carboxylic acid-functionalized polystyrene beads (d=200 µm, -COOH) were conjugated with -NH_2_ via EDC-NHS coupling using a commercial protocol [71]. In brief, 10 mg of 200 µm polystyrene beads (-COOH) were suspended in 10 mL of double-distilled water (ddH_2_O) (10 mg/mL). Sulfo-NHS and 1-(3-dimethylaminoproyl)-3-ethylcarbodiimide hydrochloride (EDC-HCl) were sequentially added to a final concentration of 0.25 mg/mL to the bead suspension and shaken for 20 min at room temperature. Then, sodium bicarbonate (NaHCO_3_, 50 mM, PH=8) and ethylenediamine (5 mg/mL) were added to the bead suspension and shaken at room temperature for 2 h. After the conjugation reaction, the beads were sterilized by three 15-minute centrifuge cycles with fresh 70% ethanol and washed with sterile PBS. Prior to use, the modified polystyrene beads (-NH_2_) were coated with 5 µg/ml Collagen IV at 37°C for at least 2 h.

#### Human TSC Culture and Trophoblast Organoid Differentiation

The hTSC line, CT27 and CT29, derived from a first-trimester placenta by Okae et al.[15], was kindly provided by Dr. Michael Soares (UMKC) and was cultured in hTSC 2D medium [DMEM/F12 supplemented with 0.1mM 2-mercaptoethanol, 0.2% FBS, 0.3% bovine serum albumin (BSA), 1% I=insulin-transferrin-selenium-ethanolamine supplement (ITS-X), 1.5 mg/ml L-ascorbic acid, 50 ng/ml hEGF, 2mM CHIR99021, 0.5mM A83-01, 1mM SB431542, 0.8 mM valproic acid and 5 mM Y27632]. Cells were dissociated into single cells by TrypLE, washed with DMEM/F12 culture medium and pelleted. The cell pellet was resuspended in hTSC 2D medium and 5×10^4^ cells were incubated with 0.5 mg collagen IV pre-coated beads (approximately 40 beads) in 3 ml hTSC 2D medium supplemented with 5µM Y27623 in a round-bottom test tube at 37°C for 4 h, the tube was stirred gently for 1 minute every 30 minutes. Cells were then transferred to Ultra-Low attachment 6-well plates and cultured for additional 48 h. The surfaces of the collagen-coated beads were largely covered with cells within 24 h. On day 3, the medium was changed to trophoblast organoid medium (TOM)[16] [DMEM/F12, 1X N2 supplement, 1X B27 supplement minus vitamin A, 1.25 mM N-acetyl-L-cysteine, 1% GlutaMAX, (TOM basal medium)], supplemented with 500 nM A83-01 and 1.5mM CHIR99021 to induce trophoblast organoid formation. On day 5, the culture medium was changed to TOM2 (TOM basal medium supplemented with 500 nM A83-01, 1.5mM CHIR99021, 80 ng/ml human R-spondin1, 50 ng/ml hEGF, 100 ng/ml hFGF2, 50 ng/ml hHGF, 2mM Y-27632). The medium was replaced every 3-4 days until day 10.

#### Generation of EVT-like cells from trophoblast organoids

On day 7 of differentiation, 4-5 STB trophoblast organoids, as generated above, were transferred to an Eppendorf tube with a 200-µL wide bore pipette tip. The organoids were resuspended in 20 µL Matrigel. The drop was the placed in either 34-mm TPP dishes or MatTek Corporation dishes. Matrigel was allowed to gel at 37°C for 15 min, and the drop then overlaid with 3ml EVT differentiation medium [12] (EVTM: advanced DMEM/F12, 0.1 mM 2-mercaptoethanol, 0.3% BSA, 1% ITS-X supplement, 100 ng ml−1 NRG1, 7.5 μM A83-01 and 4 % knockout serum replacement) under an atmosphere of 5% (v/v) CO_2_/air at 37°C. The medium was changed on day 11 (4 days after Matrigel embedding) and organoids collected on day 14.

#### Whole Mount Immunofluorescence Staining and Confocal Microscopy

Organoids were fixed with 4% (v/v) paraformaldehyde (PFA) in PBS for 30 min and permeabilized in 0.4% Triton X-100/PBS for 30 min. Organoids were placed in 10% (v/v) donkey serum/5% (w/v) BSA /0.1% Triton X-100/ in PBS as a blocking reagent for 2 h. Cells were then incubated with primary antibodies (Table 1) at 4 °C for 48 h. DAPI and secondary antibody staining was performed with either Alexa Fluor 555-, 647-, or 488-labeled detection reagents at a 1:300 dilution at 4°C overnight. Samples were then mounted in mounting media and images captured under a Leica TCS SP8 confocal laser scanning microscope. For live cell imaging, organoids were grown in 35mm dishes with coverslips and, just prior to imaging, incubated for 1h at 37°C in FITC labeled HLA-G antibody at 1:100. Organoids were then washed twice in DMEM/F12, transferred to EVTM and immediately imaged under a Leica TCS SP8 confocal laser scanning microscope.

#### H&E Staining and Immunohistochemistry

Organoids were fixed in 10 % (v/v) neutral buffered formalin. Paraffin-embedded organoids were sectioned and then stained with hematoxylin-eosin (H&E) or processed for immunofluorescence staining and immunohistochemistry. For immunofluorescence staining, slides were placed in boiling citric acid buffer (pH 6.0), non-specific antigens blocked in 5% (v/v) donkey serum/5 % (w/v) BSA in PBS for 1 h, and incubated with primary antibodies (Table 1) at 4°C overnight. The sections were then incubated with fluorescent-conjugated secondary antibodies for 1 h at room temperature. Images were captured under a Zeiss Axiovert 200M with a Leica DFC290 color camera. For immunohistochemistry, sections were rehydrated through a xylene substitute and graded alcohol series. Antigen retrieval was performed as previously described [72]. Immunoperoxidase staining was performed by using a Vectastain Elite ABC-HRP kit, and images captured with a Leica DM 5500B upright microscope with a color digital camera.

#### Transmission electron microscopy

The organoids were fixed in 0.1% (w/v) glutaraldehyde/4 % formaldehyde (v/v) in 0.1 M sodium cacodylate buffer pH 7.2 at room temperature for 30 min and post-fixed with 1 % (w/v) osmium tetroxide. Specimens were dehydrated through a graded ethanol series, placed in acetone and infiltrated with Epon by means of a Pelco Microwave (Ted Pella, CA). Thin sections (85 nm) were mounted on formvar/carbon-coated slot grids and post-stained with 2 % uranyl acetate and Sato’s triple lead stain. Sections were examined on a JEOL 1400 transmission electron microscope at 80kV.

#### Scanning electron microscopy

Samples were collected and processed for scanning electron microscopy (SEM). Unless otherwise stated, all reagents were purchased from Electron Microscopy Sciences and all specimen preparation performed at the Electron Microscopy Core Facility, University of Missouri. The samples were fixed in 2 % paraformaldehyde, 2 % glutaraldehyde in 100 mM sodium cacodylate buffer pH 7.35. Fixed organoids were plated overnight on coverslips pre-coated with medium to ensure adhesion and rinsed with 100 mM sodium cacodylate buffer, pH 7.35 containing 130 mM sucrose. Secondary fixation was performed in 1 % osmium tetroxide in cacodylate buffer by using a Pelco Biowave (Ted Pella, Inc.) operated at 100 Watts for 1 min. Specimens were next incubated at 4°C for 1 h, then rinsed with cacodylate buffer and finally with distilled water. A graded dehydration series (per exchange, 100 Watts for 40 s) was performed with ethanol in the Pelco Biowave,. Samples were dried by using the Tousimis Autosamdri 815 (Tousimis, Rockville, MD) and samples were sputter coated with 5nm of platinum using the EMS 150T-ES Sputter Coater. Images were acquired with a FEI Quanta 600F scanning electron microscope (FEI, Hillsboro, OR).

#### Immunoassays

Organoid culture medium was collected on days 1, 3, 5 8, and 10. Human CG (hCG; human chorionic gonadotrophin), progesterone (P4) and PGF (placental growth factor) concentrations were measured with solid-phase sandwich hCG ELISA kits, P4 ELISA kits and PGF human ELISA kits, respectively, by following the manufacturer recommended protocols. Samples were collected from three independent experiments for each time point and were run in duplicate.

#### Real-Time qPCR

Total RNA was extracted with a RNeasy Mini kit, and RNA treated with a TURBO-DNA-free kit to remove genomic DNA. RNA was reverse-transcribed with a cDNA synthesis kit. Primers were synthesized by Integrated DNA Technologies (IDT). Primer sequences are provided in Table 2. Real-time qPCR was performed with SYBR Green PCR on a CFX Connect Real-Time PCR System (BIO-RAD). GAPDH was used as the endogenous control (reference gene). Thermal cycling conditions were as follows: 95°C for 5 min, followed by 45 cycles of: denaturation at 95 °C for 10 sec, annealing at 56 °C for 10 sec and extension at 72 °C for 10 sec. CT values were normalized to the endogenous control (GAPDH) and fold-change values calculated relative to hTSC grown in 2D by the 2^−ΔΔCT^ procedure described elsewhere [73]. Results were reported as relative expression levels compared to the individual control sample in each assay.

**Tab 2.**
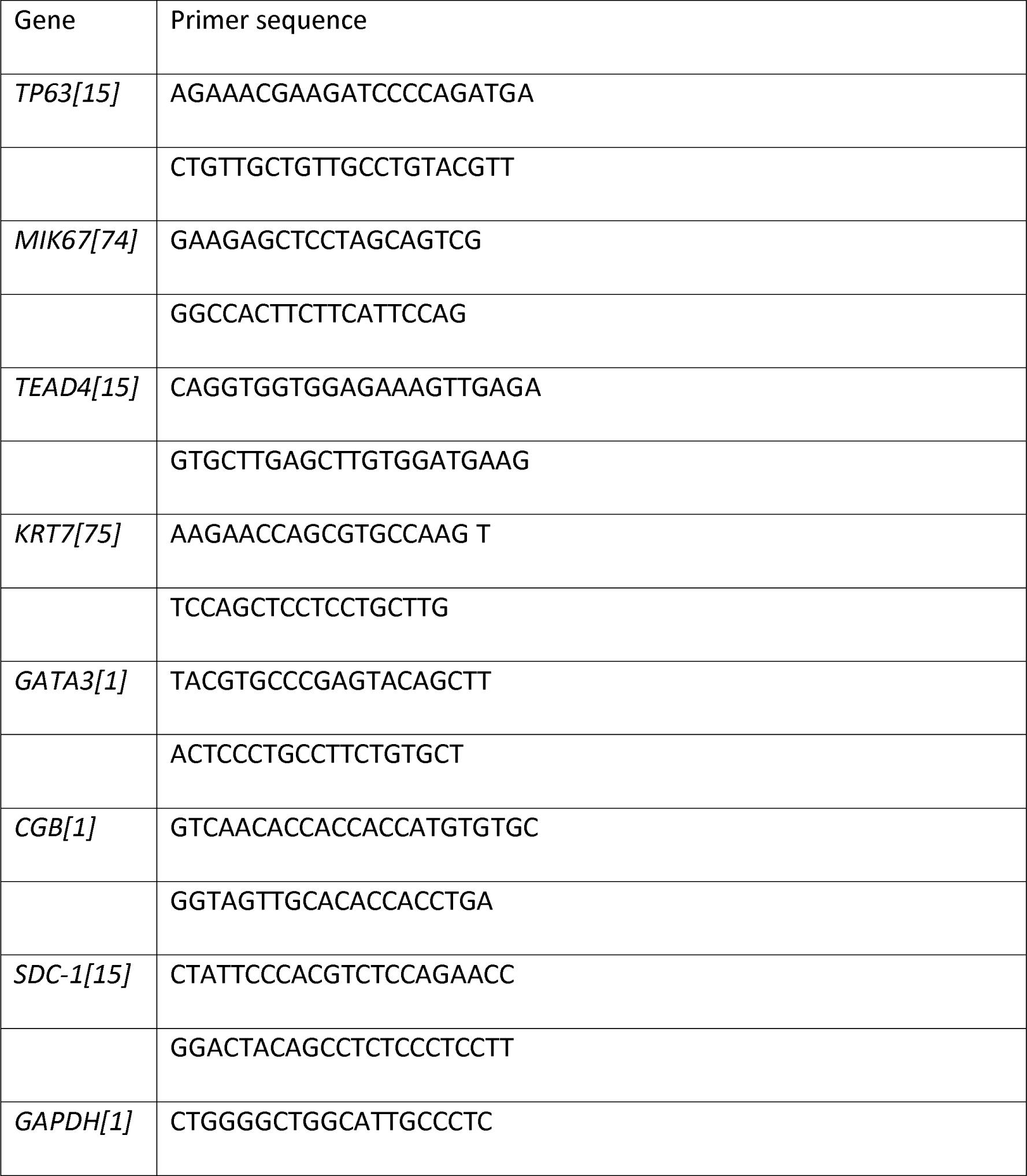
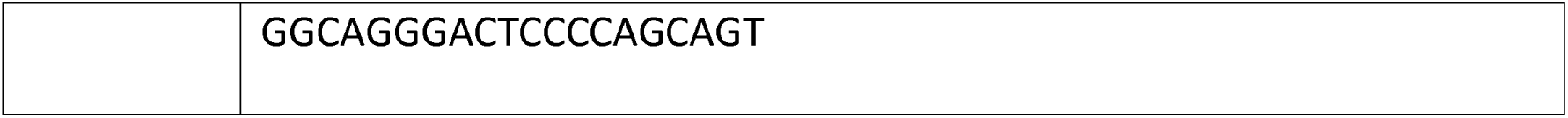
Primers used for qPCR.

### Quantification and statistical analysis

Statistical analyses were performed with GraphPad Prism 5 software. Comparisons of two groups on immunoassays were made with the Student’s t test. Values of P ≤ 0.05 were considered consistent with their mean values being different.

## Data and code availability

All data reported in this paper will be shared by the lead contact upon request.

This paper does not report original code.

Any additional information required to reanalyze the data reported in this paper is available from the lead contact upon request.

## Acknowledgements

We thank Michael J. Soares for supplying the hTSC lines, and Hiroaki Okae and Takahiro Arima of Tohoku University Graduate School of Medicine, Japan, for sharing permission. We are also thankful to Dr. Chuyu Hayashi for cell culture support. We thank the Advanced Light Microscopy and Electron Microscopy core facilities (University of Missouri) for technical support, services and training, including the microscopy core (Alexander Jurkevich and Frank Baker) and the electron microscopy core (Deana G Grant and David Stalla).

## References

1. Yang, Y., et al., Heightened potency of human pluripotent stem cell lines created by transient BMP4 exposure. Proceedings of the National Academy of Sciences, 2015. 112(18): p. E2337–E2346.

2. Jang, Y.J., et al., Induction of human trophoblast stem-like cells from primed pluripotent stem cells. Proc Natl Acad Sci U S A, 2022. 119(20): p. e2115709119.

3. Cindrova-Davies, T. and A.N. Sferruzzi-Perri. Human placental development and function. in Seminars in Cell & Developmental Biology. 2022. Elsevier.

4. Roberts, R.M. and S.J. Fisher, Trophoblast stem cells. Biol Reprod, 2011. 84(3): p. 412–21.

5. Khan, T., et al., Single Nucleus RNA Sequence (snRNAseq) Analysis of the Spectrum of Trophoblast Lineages Generated From Human Pluripotent Stem Cells in vitro. Front Cell Dev Biol, 2021. 9: p. 695248.

6. Genbacev, O., S.A. Schubach, and R.K. Miller, Villous culture of first trimester human placenta--model to study extravillous trophoblast (EVT) differentiation. Placenta, 1992. 13(5): p. 439–61.

7. Pijnenborg, R., et al., Trophoblastic invasion of human decidua from 8 to 18 weeks of pregnancy. Placenta, 1980. 1(1): p. 3–19.

8. Aplin, J.D., Developmental cell biology of human villous trophoblast: current research problems. Int J Dev Biol, 2010. 54(2-3): p. 323–9.

9. Evain-Brion, D. and A. Malassine, Human placenta as an endocrine organ. Growth Horm IGF Res, 2003. 13 Suppl A: p. S34–7.

10. Johnson, M.R., et al., The role of trophoblast dysfunction in the aetiology of miscarriage. Br J Obstet Gynaecol, 1993. 100(4): p. 353–9.

11. Huppertz, B., The Critical Role of Abnormal Trophoblast Development in the Etiology of Preeclampsia. Curr Pharm Biotechnol, 2018. 19(10): p. 771–780.

12. Scifres, C.M. and D.M. Nelson, Intrauterine growth restriction, human placental development and trophoblast cell death. J Physiol, 2009. 587(Pt 14): p. 3453–8.

13. Burton, G.J. and E. Jauniaux, What is the placenta? Am J Obstet Gynecol, 2015. 213(4 Suppl): p. S6.e1, S6-8.

14. Kliman, H.J., et al., Purification, characterization, and in vitro differentiation of cytotrophoblasts from human term placentae. Endocrinology, 1986. 118(4): p. 1567–82.

15. Okae, H., et al., Derivation of human trophoblast stem cells. Cell stem cell, 2018. 22(1): p. 50–63. e6.

16. Turco, M.Y., et al., Trophoblast organoids as a model for maternal–fetal interactions during human placentation. Nature, 2018. 564(7735): p. 263–267.

17. Haider, S., et al., Self-renewing trophoblast organoids recapitulate the developmental program of the early human placenta. Stem cell reports, 2018. 11(2): p. 537–551.

18. Sato, M., et al., Three-dimensional human placenta-like bud synthesized from induced pluripotent stem cells. Sci Rep, 2021. 11(1): p. 14167.

19. Cui, K., et al., Establishment of Trophoblast-Like Tissue Model from Human Pluripotent Stem Cells in Three-Dimensional Culture System. Adv Sci (Weinh), 2022. 9(3): p. e2100031.

20. Jones, C.J. and H. Fox, Ultrastructure of the normal human placenta. Electron Microsc Rev, 1991. 4(1): p. 129–78.

21. Castillo, D.D.L.R., et al., Atypical cristae morphology of human syncytiotrophoblast mitochondria: role for complex V. Journal of Biological Chemistry, 2011. 286(27): p. 23911–23919.

22. Smith, C.A., H.D. Moore, and J.P. Hearn, The ultrastructure of early implantation in the marmoset monkey (Callithrix jacchus). Anat Embryol (Berl), 1987. 175(3): p. 399–410.

23. Palmer, M.E., A.L. Watson, and G.J. Burton, Morphological analysis of degeneration and regeneration of syncytiotrophoblast in first trimester placental villi during organ culture. Hum Reprod, 1997. 12(2): p. 379–82.

24. Simán, C.M., et al., The functional regeneration of syncytiotrophoblast in cultured explants of term placenta. Am J Physiol Regul Integr Comp Physiol, 2001. 280(4): p. R1116–22.

25. Sheridan, M.A., et al., Characterization of primary models of human trophoblast. Development, 2021. 148(21).

26. Holland, O., et al., Review: Placental mitochondrial function and structure in gestational disorders. Placenta, 2017. 54: p. 2–9.

27. Martínez, F., M. Kiriakidou, and J.F. Strauss III, Structural and functional changes in mitochondria associated with trophoblast differentiation: methods to isolate enriched preparations of syncytiotrophoblast mitochondria. Endocrinology, 1997. 138(5): p. 2172–2183.

28. Kolahi, K.S., A.M. Valent, and K.L. Thornburg, Cytotrophoblast, Not Syncytiotrophoblast, Dominates Glycolysis and Oxidative Phosphorylation in Human Term Placenta. Sci Rep, 2017. 7: p. 42941.

29. Mizushima, N., T. Yoshimori, and B. Levine, Methods in mammalian autophagy research. Cell, 2010. 140(3): p. 313–26.

30. Bastida-Ruiz, D., et al., The fine-tuning of endoplasmic reticulum stress response and autophagy activation during trophoblast syncytialization. Cell Death Dis, 2019. 10(9): p. 651.

31. Ockleford, C.D., et al., Structure and function of the nematosome. J Cell Sci, 1987. 87 (Pt 1): p. 27–44.

32. Sheridan, M.A., et al., Establishment and differentiation of long-term trophoblast organoid cultures from the human placenta. Nature Protocols, 2020. 15(10): p. 3441–3463.

33. Aplin, J.D., et al., Development of Cytotrophoblast Columns from Explanted First-Trimester Human Placental Villi: Role of Fibronectin and Integrin α5β11. Biology of Reproduction, 1999. 60(4): p. 828–838.

34. Biadasiewicz, K., et al., Extravillous Trophoblast-Associated ADAM12 Exerts Pro-Invasive Properties, Including Induction of Integrin Beta 1-Mediated Cellular Spreading1. Biology of Reproduction, 2014. 90(5).

35. West, R.C., et al., Dynamics of trophoblast differentiation in peri-implantation–stage human embryos. Proceedings of the National Academy of Sciences, 2019. 116(45): p. 22635–22644.

36. Lee, C.Q.E., et al., Integrin α2 marks a niche of trophoblast progenitor cells in first trimester human placenta. Development, 2018. 145(16).

37. Shirendeb, U., et al., Human papillomavirus infection and its possible correlation with p63 expression in cervical cancer in Japan, Mongolia, and Myanmar. Acta Histochem Cytochem, 2009. 42(6): p. 181–90.

38. Hooghe, B., et al., ConTra: a promoter alignment analysis tool for identification of transcription factor binding sites across species. Nucleic Acids Res, 2008. 36(Web Server issue): p. W128–32.

39. Guttmacher, A.E., Y.T. Maddox, and C.Y. Spong, The Human Placenta Project: placental structure, development, and function in real time. Placenta, 2014. 35(5): p. 303–4.

40. Karvas, R.M., et al., Stem-cell-derived trophoblast organoids model human placental development and susceptibility to emerging pathogens. Cell Stem Cell, 2022. 29(5): p. 810–825. e8.

41. Marinić, M., S. Rana, and V.J. Lynch, Derivation of endometrial gland organoids from term placenta. Placenta, 2020. 101: p. 75–79.

42. Schust, D.J., et al., The Immunology of Syncytialized Trophoblast. Int J Mol Sci, 2021. 22(4).

43. Coleman, S.J., et al., Syncytial nuclear aggregates in normal placenta show increased nuclear condensation, but apoptosis and cytoskeletal redistribution are uncommon. Placenta, 2013. 34(5): p. 449–55.

44. Göhner, C., T. Plösch, and M.M. Faas, Immune-modulatory effects of syncytiotrophoblast extracellular vesicles in pregnancy and preeclampsia. Placenta, 2017. 60: p. S41–S51.

45. Gorrill, M.J., et al., Initial experience with extended culture and blastocyst transfer of cryopreserved embryos. American journal of obstetrics and gynecology, 1999. 180(6 Pt 1): p. 1472–1474.

46. VanPutte, C.L., et al., Seeley’s anatomy & physiology. 2019: University of Iowa.

47. Simon, B.H., H.Y. Ando, and P.K. Gupta, Circulation time and body distribution of 14C-labeled amino-modified polystyrene nanoparticles in mice. J Pharm Sci, 1995. 84(10): p. 1249–53.

48. Schwarzbauer, J., Basement membrane: Putting up the barriers. Current Biology, 1999. 9(7): p. R242–R244.

49. Oefner, C.M., et al., Collagen type IV at the fetal-maternal interface. Placenta, 2015. 36(1): p. 59–68.

50. Jones, C.J. and H. Fox, Ultrastructure of the normal human placenta. Electron microscopy reviews, 1991. 4(1): p. 129–178.

51. Boyd, J. and A. Hughes, Observations on human chorionic villi using the electron microscope. Journal of Anatomy, 1954. 88(Pt 3): p. 356.

52. Terzakis, J.A., The ultrastructure of normal human first trimester placenta. Journal of Ultrastructure Research, 1963. 9(3-4): p. 268–284.

53. Pierce Jr, G. and A. Midgley Jr, The origin and function of human syncytiotrophoblastic giant cells. The American Journal of Pathology, 1963. 43(2): p. 153.

54. Pierce Jr, G., A. Midgley Jr, and B. TF, AN ULTRASTRUCTURAL STUDY OF DIFFERENTIATION AND MATURATION OF TROPHOBLAST OF THE MONKEY. Laboratory Investigation; a Journal of Technical Methods and Pathology, 1964. 13: p. 451–464.

55. De Falco, S., The discovery of placenta growth factor and its biological activity. Exp Mol Med, 2012. 44(1): p. 1–9.

56. Selvaraj, S.K., et al., Mechanism of monocyte activation and expression of proinflammatory cytochemokines by placenta growth factor. Blood, 2003. 102(4): p. 1515–1524.

57. Oura, H., et al., A critical role of placental growth factor in the induction of inflammation and edema formation. Blood, The Journal of the American Society of Hematology, 2003. 101(2): p. 560–567.

58. Tudisco, L., et al., Hypoxia activates placental growth factor expression in lymphatic endothelial cells. Oncotarget, 2017. 8(20): p. 32873–32883.

59. Al-Zuhair, A., et al., Loss and regeneration of the microvilli of human placental syncytiotrophoblast. Archives of gynecology, 1987. 240(3): p. 147–151.

60. Enders, A.C., Trophoblast differentiation during the transition from trophoblastic plate to lacunar stage of implantation in the rhesus monkey and human. Am J Anat, 1989. 186(1): p. 85–98.

61. Lange, K., Fundamental role of microvilli in the main functions of differentiated cells: Outline of an universal regulating and signaling system at the cell periphery. J Cell Physiol, 2011. 226(4): p. 896–927.

62. Guillomot, M., Cellular interactions during implantation in domestic ruminants. Journal of Reproduction and Fertility-Supplements only, 1995(49): p. 39–52.

63. Gonda, M.A. and Y.C. Hsu, Correlative scanning electron, transmission electron, and light microscopic studies of the in vitro development of mouse embryos on a plastic substrate at the implantation stage. J Embryol Exp Morphol, 1980. 56: p. 23–39.

64. Highison, G.J. and F.D. Tibbitts, Ultrasonic microdissection of immature intermediate human placental villi as studied by scanning electron microscopy. Scanning Electron Microscopy, 1986. 1986(2): p. 38.

65. Carter, A.M., A.C. Enders, and R. Pijnenborg, The role of invasive trophoblast in implantation and placentation of primates. Philos Trans R Soc Lond B Biol Sci, 2015. 370(1663): p. 20140070.

66. Zhuang, B., J. Shang, and Y. Yao, HLA-G: An Important Mediator of Maternal-Fetal Immune-Tolerance. Front Immunol, 2021. 12: p. 744324.

67. Banet, N., et al., GATA-3 expression in trophoblastic tissues: an immunohistochemical study of 445 cases, including diagnostic utility. Am J Surg Pathol, 2015. 39(1): p. 101–8.

68. Lee, Y., et al., A unifying concept of trophoblastic differentiation and malignancy defined by biomarker expression. Hum Pathol, 2007. 38(7): p. 1003–1013.

69. Li, Y., et al., p63 inhibits extravillous trophoblast migration and maintains cells in a cytotrophoblast stem cell-like state. Am J Pathol, 2014. 184(12): p. 3332–43.

70. Bulmer, J.N., L. Morrison, and P.M. Johnson, Expression of the proliferation markers Ki67 and transferrin receptor by human trophoblast populations. J Reprod Immunol, 1988. 14(3): p. 291–302.

71. Coelho, N.M., et al., Arrangement of type IV collagen on NH₂ and COOH functionalized surfaces. Biotechnol Bioeng, 2011. 108(12): p. 3009–18.

72. Karvas, R.M., et al., Use of a human embryonic stem cell model to discover GABRP, WFDC2, VTCN1 and ACTC1 as markers of early first trimester human trophoblast. Molecular human reproduction, 2020. 26(6): p. 425–440.

73. Schmittgen, T.D. and K.J. Livak, Analyzing real-time PCR data by the comparative C T method. Nature protocols, 2008. 3(6): p. 1101.

74. Hachim, M.Y., et al., Derangement of cell cycle markers in peripheral blood mononuclear cells of asthmatic patients as a reliable biomarker for asthma control. Scientific reports, 2021. 11(1): p. 1–24.

75. Amita, M., et al., Complete and unidirectional conversion of human embryonic stem cells to trophoblast by BMP4. Proceedings of the National Academy of Sciences, 2013. 110(13): p. E1212–E1221.

